# Role of glycine-*N*-methyl transferase in human and murine nonalcoholic steatohepatitis

**DOI:** 10.1101/2021.12.03.470998

**Authors:** Connor R. Quinn, Mario C. Rico, Carmen Merali, Salim Merali

## Abstract

Non-alcoholic fatty liver disease (NAFLD) has become one of the most prominent forms of chronic liver disease worldwide, mirroring the obesity epidemic. NAFLD is the number one cause of chronic liver disease worldwide, with 25% of these patients developing nonalcoholic steatohepatitis (NASH). This significantly increases the risk of cirrhosis and decompensated liver failure. Past studies in rodent models have shown that the knockout of glycine-*N*-methyltransferase (GNMT) is associated with steatosis, fibrosis, and hepatocellular carcinoma. However, the attenuation of GNMT in subjects with NASH and the molecular basis for its impact on the disease process have yet to be elucidated. To address this knowledge gap, we show the reduction of GNMT protein levels in the liver of NASH subjects compared to healthy controls. To gain insight into the impact of decreased GNMT in the disease process, we performed global label-free proteome studies on the livers from a murine Western diet-based model of NASH. We identified the differential expression of essential proteins involved in hallmark NASH pathogenesis, including lipid metabolism, inflammation, and fibrosis. Significantly, the downregulation of GNMT, the prominent regulator of *S*-adenosylmethionine (AdoMet), was identified as a contributing factor to these networks, increasing fourfold in AdoMet levels. AdoMet is an essential metabolite for transmethylation reactions and a substrate for polyamine synthesis, and its levels can impact polyamine flux and generate oxidative stress. Therefore, we performed targeted proteome and metabolomics studies to show a decrease in GNMT transmethylation, an increase in flux through the polyamine pathway, and increased oxidative stress.

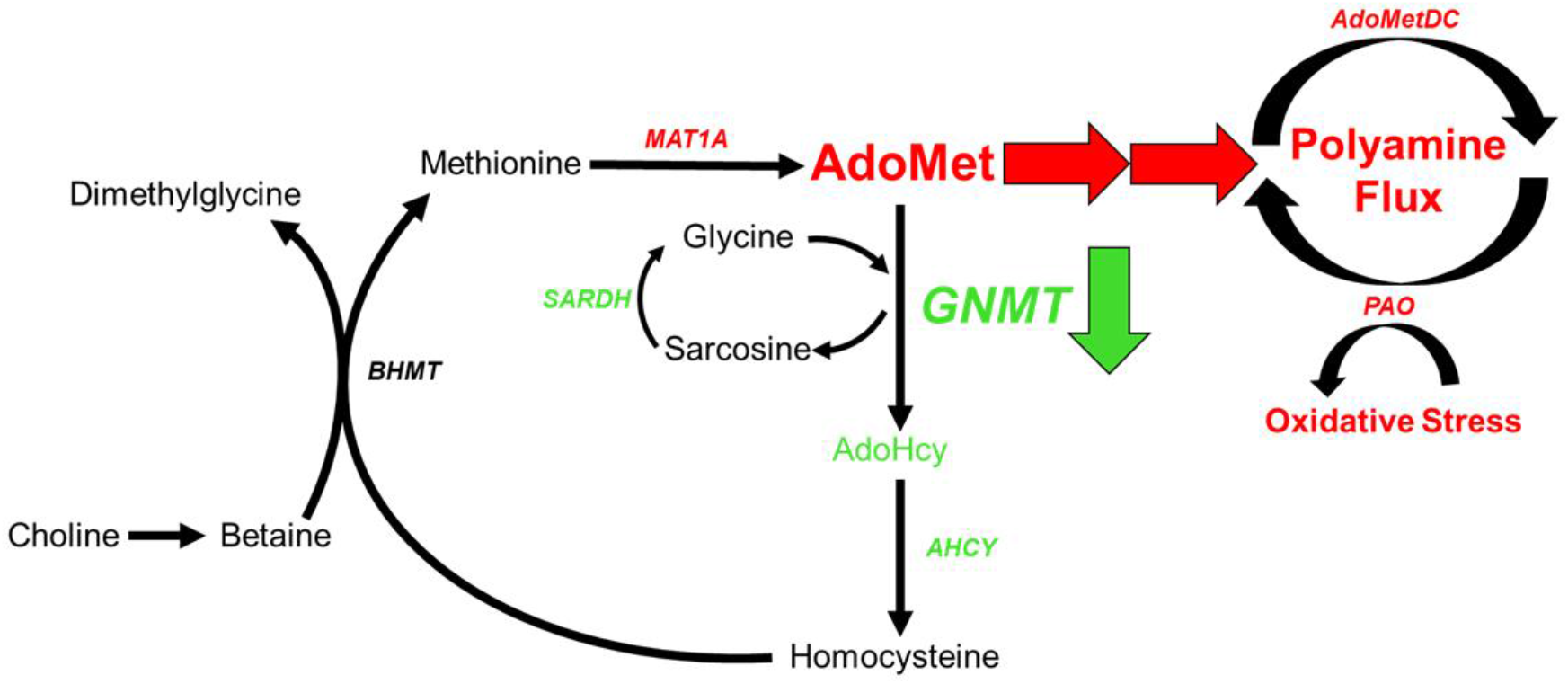

## Introduction

Nonalcoholic fatty liver disease (NAFLD) is an excess buildup of fat in the liver and is the leading cause of chronic liver disease worldwide. Currently, it is estimated 25% of the human population has NAFLD, and of these patients, 25% will progress to the more severe debilitating disease nonalcoholic steatohepatitis (NASH) (Younossi et al., 2018; Younossi et al., 2019). NASH is characterized by hepatocellular damage involving steatosis, inflammation, and fibrosis of the liver. It is a major risk factor for cirrhosis and hepatocellular carcinoma, and it significantly increases the risk of death due to liver-related causes (Younossi *et al*., 2018; Younossi *et al*., 2019). Currently, there are no FDA-approved treatments on the market for treating NASH and current therapies are limited to lifestyle interventions (calorie restriction, exercise) and treating comorbidities (insulin resistance, obesity). Due to low patient compliance and the growing obesity epidemic, the number of patients diagnosed with NAFLD and NASH continues to grow (Younossi and Henry, 2019).

The pathogenesis of NASH has been described by the “multiple hit theory,” where several metabolic insults, including increased de novo lipogenesis, mitochondrial dysfunction, inflammation, and oxidative stress, simultaneously lead to the disease state (Buzzetti et al., 2016). The molecular mechanisms driving these insults are still not well understood. Glycine-*N*-methyl transferase (GNMT) is the most abundant methyltransferase in the liver, accounting for over 1% of all cytosolic protein in hepatocytes (Mato et al., 2013; Mudd et al., 2007). It catalyzes the transfer of a methyl group from *S*-adenosylmethionine (AdoMet), the universal methyl donor, to glycine to form sarcosine. Because of its high abundance, GNMT plays a crucial role in controlling AdoMet metabolism and methyltransferase reactions involving AdoMet. 75% of all transmethylation reactions occur in the liver, making AdoMet a vital component of liver metabolism and regulation (Mudd *et al*., 2007). The importance of GNMT in the progression of NAFLD has been demonstrated in several studies. GNMT KO animals spontaneously develop steatosis around 3 months of age and inflammation and fibrosis of the liver at 8 months (Hughey et al., 2018; Martínez-Chantar et al., 2008). These studies indicated GNMT KO results in a significant increase in liver AdoMet that is directly responsible for steatosis and fibrosis. It was shown elevated AdoMet could lead to hypermethylation of DNA and modulation of lipid metabolism through phosphatidylcholine synthesis. Furthermore, multiple genetic analyses of human patients have found GNMT gene expression is repressed in both NASH and hepatocellular carcinoma patients (Moylan et al., 2014; Ryaboshapkina and Hammar, 2017; Teufel et al., 2016). Despite this, the GNMT protein downregulation and its impact on the disease process is still not fully understood.

In addition to transmethylation reactions, AdoMet is a required precursor for polyamine catabolism. AdoMet can donate its aminopropyl group to putrescine, the basic polyamine building block, to produce spermine and spermidine. Polyamines are polycations that play a role in multiple cellular processes, including cellular proliferation, differentiation, and death. The role of polyamine metabolism has been indicated in several disease states, including cancer and obesity (Casero et al., 2018; Jell et al., 2007). Previously, we have demonstrated the role of polyamine flux and acetyl-CoA utilization in lipid accumulation in adipose tissue (Jell *et al*., 2007; Kramer et al., 2008; Liu et al., 2014). To date, the role of polyamine metabolism has not been studied in the pathophysiology of NASH.

In this study, we show the reduction of liver GNMT protein in subjects with NASH compared to healthy controls. To further assess the effects of GNMT downregulation in NASH, we used a diet-induced NASH animal model to show that GNMT downregulation leads to a significant increase in AdoMet levels. Using targeted proteomics and metabolomics, we also show GNMT downregulation results in reduced transmethylation activity and increased flux into polyamine metabolism. Activation of polyamine flux results in increased oxidative stress. This study provides key insights into metabolic changes in the liver contributing to a cytotoxic environment and NASH pathogenesis.

## Results

### GNMT is downregulated in human NASH

Multiple studies have shown GNMT mRNA levels are decreased in NASH subjects, and there is a negative correlation between the severity of liver damage and GNMT expression (Moylan *et al*., 2014; Ryaboshapkina and Hammar, 2017). We developed and validated a targeted mass spectrometry method using selected reaction monitoring (SRM) to identify and quantitate liver GNMT in NASH and healthy subjects. The method had a coefficient of variation below 15% and 5% for technical and SRM replicates, respectively and the peak area response was linear from 10-200µg of total protein (R^2^=0.997) (Suppl. Fig 1 and 2). The control group consisted of 3 male and 2 female subjects with a mean age of 43 and a normal BMI. The NASH subjects included 2 males and 3 females with a mean age of 54. All NASH subjects had a BMI greater than 30 and were diagnosed with diabetes. We show GNMT liver protein levels were decreased threefold in NASH subjects compared to healthy subjects (Figure 1).

**Fig 1.**
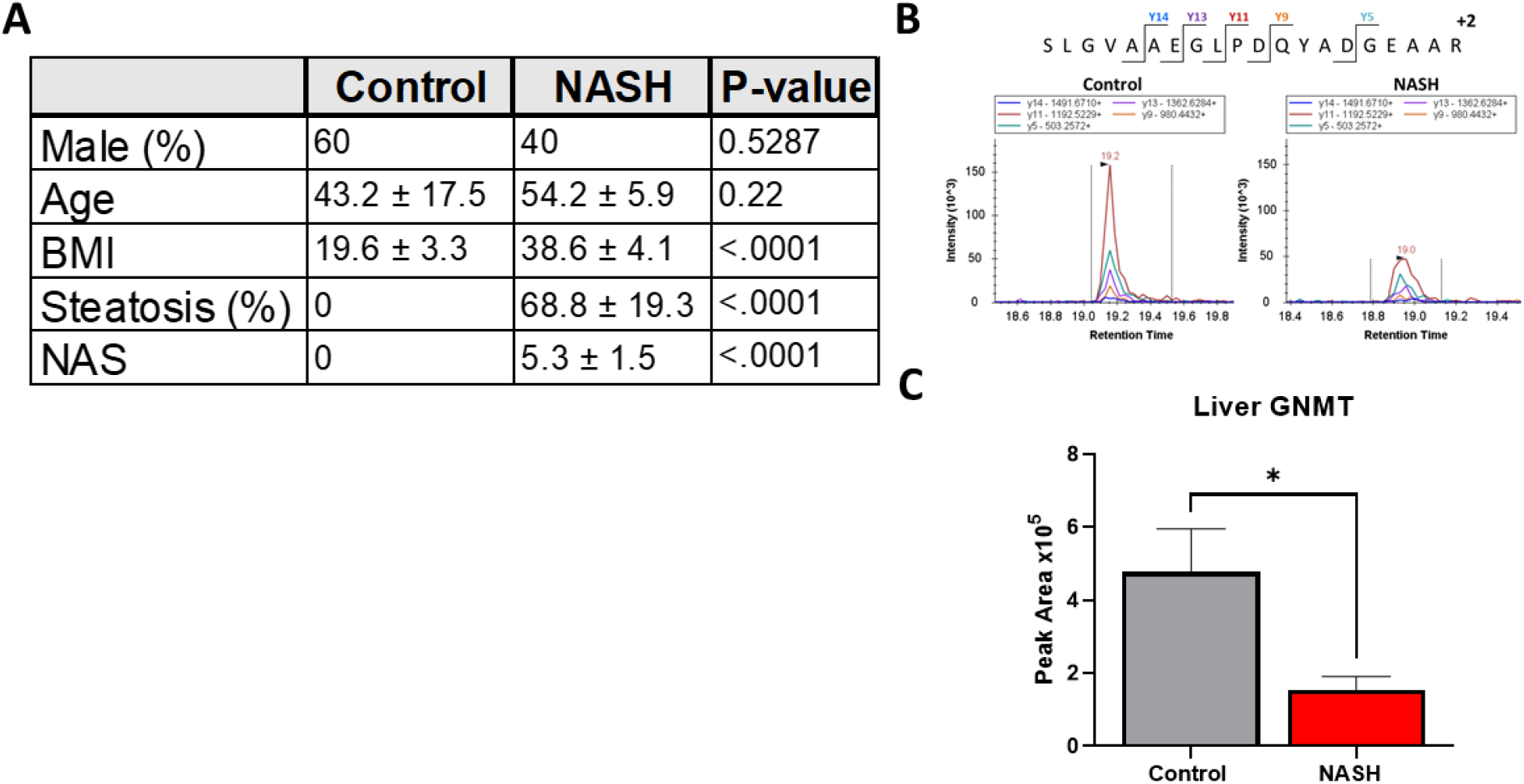
GNMT is downregulated in humanNASH liver. (A) Table displaying patient characteristics for control and NASH liver samples.(B) Representative chromatograms from selected reaction monitoring analysis (C) GNMT protein abundance in human liver samples. Data presented as mean ± SEM. (N=5 *p<.05)

### Western diet induces NASH in mice

To better understand the effect of GNMT reduction in NASH progression, we modified a diet-induced animal model for studying NASH pathophysiology. Specifically, a modified Amylin diet with high fat, fructose, and cholesterol was used to model NASH in mice. Recent studies have shown high-fat diets with high cholesterol can induce inflammation and fibrosis in rodent liver (Charlton et al., 2011; Kohli et al., 2010) and transcriptomic and metabolic analyses show these diets can accurately model human NASH and activate key pathways involved in human NASH progression (Machado et al., 2015; Teufel *et al*., 2016). C57BL-6J mice were fed a Western diet for 58 weeks to induce NASH. The Western diet animals showed a significant increase in weight compared to control animals and demonstrated insulin resistance during glucose tolerance test (Figure 2B/C). At the end of the study, the liver tissue was collected, and liver tissue structure and integrity were visualized by H&E, Mason Trichrome, and Sirius red staining (Figure 2A). Tissue damage and cell death were evident in the Western diet animals from cytoplasm clearing, cell enlargement, and immune cell infiltration. The NAFLD Activity Score (NAS) and Fibrosis score were used to confirm the diagnosis of NASH (The diagnosis and management of nonalcoholic fatty liver disease: Practice guidance from the American Association for the Study of Liver Diseases, 2018; Liang et al., 2014). The NAS scoring system is based on three parameters steatosis, lobular inflammation, and hepatic ballooning. The Western diet animals had significantly higher levels of all three parameters compared to control and an average NAS score of 6. The degree of fibrosis was assessed by Mason trichrome and Sirius red staining of collagen fibers. The Western diet animals had an average Fibrosis score of 2 (Figure 2D). This analysis verified the diagnosis of NASH with fibrosis in Western diet animals and validated our studies on NASH pathophysiology.

**Fig 2.**
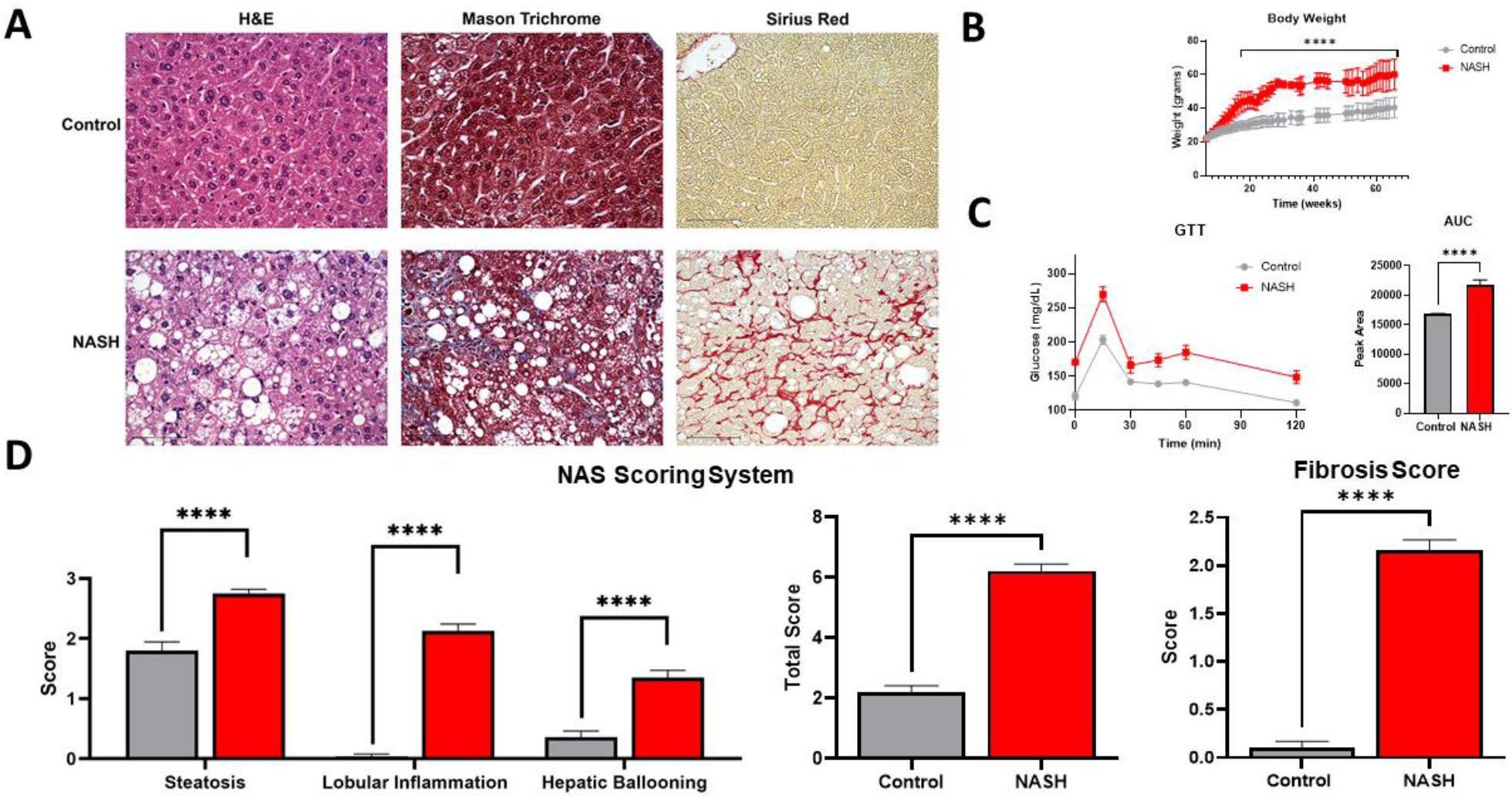
Western diet inducesNASH in mice. (A) H&E, Masson trichrome, and Sirius Red liver histology staining. Scale bar=75µm (B) Bodyweight of animals (C) Results from glucose tolerance test and area under the curve quantitation (C) The NAS scoring system was used to assess liver histology. Animals fed a western diet were diagnosed with NASH with a total average NAS score of6 and fibrosis score of 2. Data presented as mean ± SEM. (N=7 ***p<.001 ****p<.0001)

### Proteomic characterization of NASH animal model reproduces human pathophysiology

We then used global unbiased label-free proteomics to gain insight into NASH pathophysiology on the molecular level. Liver tissue from three control and three NASH animals were collected and individually processed for proteomic analysis. Our modified in-Stage Tip (iST) sample preparation process allows for minimal sample handling and fractionation to ensure high reproducibility and sensitivity (Chen et al., 2020; McBrearty et al., 2021; Molina-Franky et al., 2021; Zhang et al., 2018). From this analysis, a total of 4,783 proteins were identified and quantified with high confidence (Table S2). There were 646 differentially expressed proteins between control and NASH groups, 418 downregulated and 228 upregulated. Several of the differentially regulated proteins are involved in NASH pathophysiology, including lipid metabolism, apoptosis, fibrosis, and inflammation (Figure 3). Specifically, GNMT, platelet glycoprotein-4 (CD36), and acyl-coenzyme A thioesterase 9 (Acot9) are involved in lipid metabolism and have previously been associated with human NAFLD progression (García-Monzón et al., 2014; Martínez-Chantar *et al*., 2008; Steensels et al., 2020; Wilson et al., 2016). Collagen type 1 alpha 1 chain (COL1A1), actin alpha 2 (ACTA2), and decorin (Dcn) are three of the major markers of extracellular matrix deposition and liver fibrogenesis in humans (Friedman, 2008). Inflammatory and immunoregulatory proteins, galectin-3 (Lgals3) and interferon-induced protein with tetratricopeptide repeat 3 (Ifit3) were both upregulated in our NASH model in agreement with previous studies (Chalasani et al., 2020; Iacobini et al., 2011; Xiong et al., 2019). Currently galectin-3 inhibitors are being tested in the clinic for the treatment of NASH (Chalasani *et al*., 2020). We then used Ingenuity Pathway Analysis (IPA) software to analyze the differentially expressed proteins for overrepresented canonical pathways, regulators, and biological processes (Figure 4). IPA identified acute phase response signaling as the top altered canonical pathway, demonstrating immune response and inflammation in our animal model. Major nuclear receptor pathways were also identified as differentially regulated including liver X receptors (LXR), farnesoid X receptor (FXR), and retinoid X receptor (RXR). These nuclear receptors regulate liver metabolism and immunology and have been shown to be viable drug targets for treating NASH in the clinic (Cariello et al., 2021). Furthermore, pathways involved in fibrosis signaling and oxidative stress production were shown to be activated in our animal model. Overall, these findings correlate well with human NASH pathophysiology and validate our animal model for studying NASH on the molecular level. IPA identified proteins contributing to steatohepatitis based on previous reports and differential protein expression. In accordance with our initial human liver analysis, the downregulation of GNMT was identified as a contributing factor to steatohepatitis. Despite this, the effects of GNMT attenuation under pathophysiological conditions are still not well understood.

**Fig 3.**
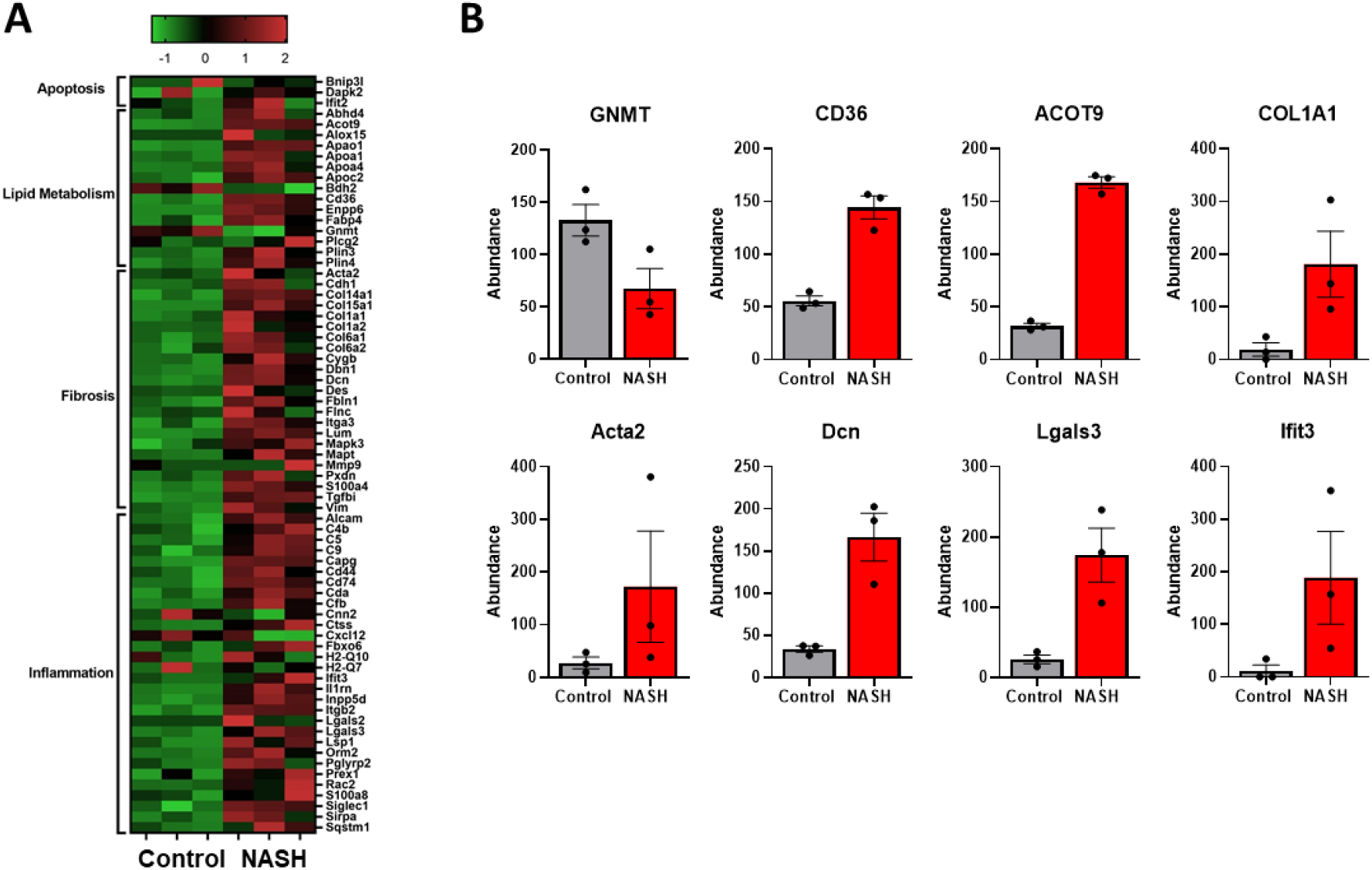
Proteomic characterization of NASH animal model reproduces human pathophysiology. (A) Heat map displaying differentially regulated proteins involved in apoptosis, lipid metabolism, fibrosis, and inflammation. (B) Individual bar graphs for markers of NASH. Glycine N-methyltransferase (GNMT), platelet glycoprotein 4 (CD36), acyl-coenzyme A thioesterase 9 (Acot9), collagen type 1 alpha 1 chain (COL1A1), actin alpha 2 (Acta2), decorin (Den), galectin-3 (Lgals3), interferon-induced proteinwith tetratricopeptide repeats 3 (lfit3).

**Fig 4.**
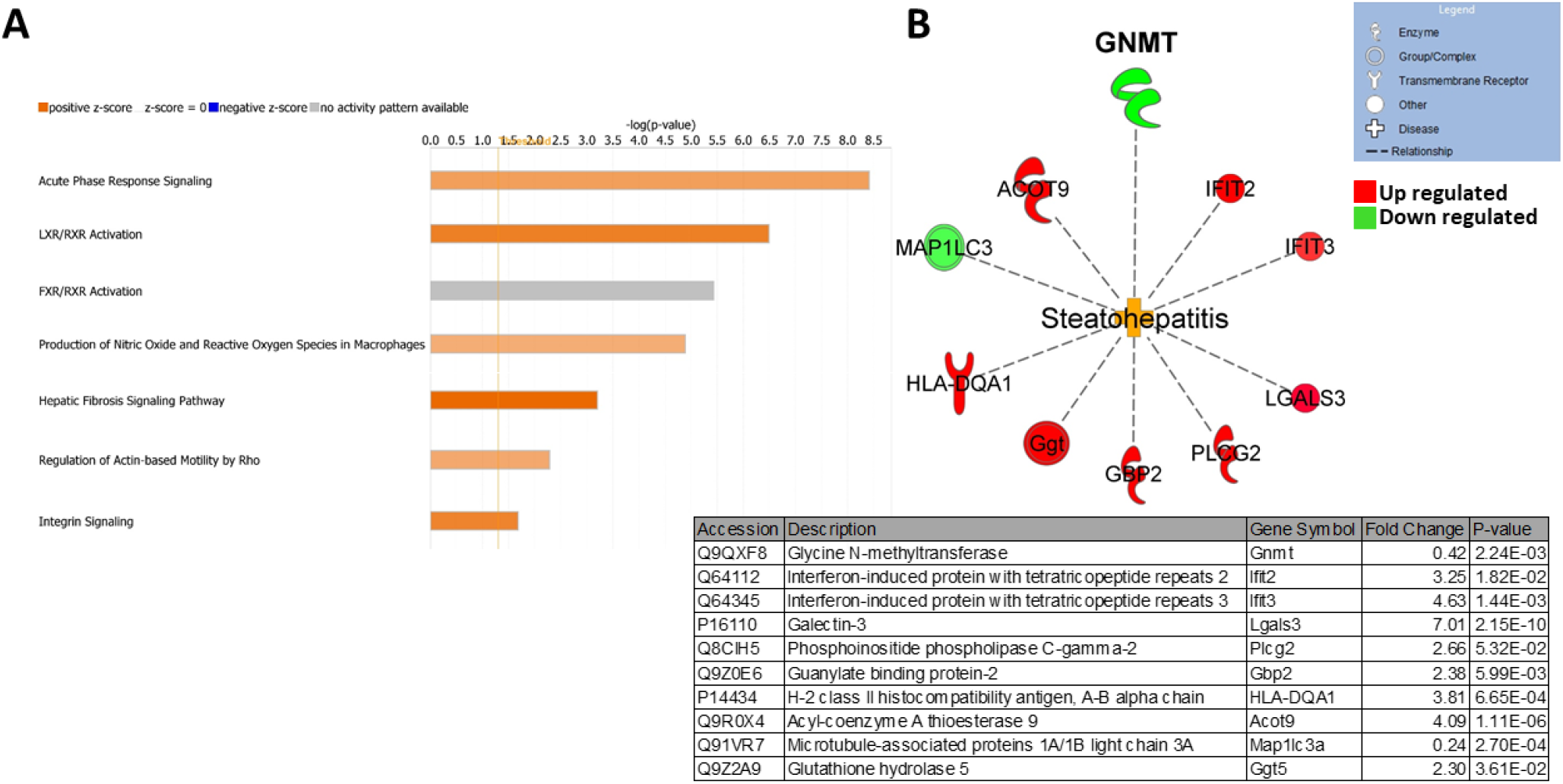
Ingenuity pathway analysis predicts activationofsteatohepatitis. (A) Top altered canonical pathways in NASH liver (B) Steatohepatitis network of differentially expressedproteins contributing to the disease state. Individual proteins are listed in the table below.

### Dysregulation of AdoMet metabolism in NASH

GNMT is highly abundant in hepatocytes and is the main regulator of AdoMet levels in the liver. Next, using a new group of control and NASH mice (n=4), we wanted to confirm the down regulation of GNMT and further investigate the effect on AdoMet regulation. A targeted mass spectrometry method was developed and validated to quantify proteins involved in AdoMet regulation. We found both GNMT and sarcosine dehydrogenase (SARDH) were downregulated in NASH. Together, GNMT and SARDH form a futile cycle in the methylation and demethylation of glycine and sarcosine to control AdoMet concentration. Furthermore, we found adenosylhomocysteinase (AHCY), an enzyme downstream of GNMT in the transmethylation pathway was downregulated and AdoMet biosynthetic enzyme, methionine adeonsyltransferase 1A (MAT1A) was significantly upregulated in NASH (Figure 4A). We then measured the levels of AdoMet and S-adenosylhomocysteine (AdoHcy) in the liver. There was a fourfold increase in AdoMet levels and approximately 30% reduction in AdoHcy in NASH (Figure 4B). This resulted in a significant increase in the AdoMet/AdoHcy ratio. These results show AdoMet regulation is disrupted in NASH inducing significantly higher concentrations of AdoMet.

### Polyamine metabolism is activated in NASH causing a flux and oxidative stress

In addition to transmethylation, AdoMet is a required substrate for polyamine synthesis and AdoMet levels can impact polyamine flux. Based on the increased levels of AdoMet we predicted to find activation of polyamine metabolism. Targeted mass spectrometry analysis found a significant increase in both rate-limiting biosynthetic enzyme S-adenosylmethionine decarboxylase (AdoMetDC) and catabolic enzyme spermidine/spermine-N1-acetyltransferase (SSAT1) (Figure 5A). Quantification of key polyamine metabolites revealed a significant increase in the polyamine building block putrescine. From our analysis there was no change in spermidine and spermine levels, but there was a significant increase in 5’-methylthioadenosine (MTA), the side product of spermidine and spermine synthesis (Figure 5B). Polyamine catabolism involves acetylation of spermine and spermidine by SSAT1 and acetyl CoA. The acetylated products can then be exported from the cell or oxidized to produce putrescine, hydrogen peroxide, and reactive aldehyde species. We found the activity of catabolic enzyme polyamine oxidase (PAO) was significantly increased in NASH (Figure 6A). Together, these findings demonstrate an increased flux into spermidine and spermine synthesis, and rapid export or degradation back to putrescine to maintain equilibrium. High rates of polyamine metabolism and polyamine oxidase activity can create high levels of oxidative stress through the production of hydrogen peroxide and reactive aldehyde species. The degree of oxidative damage in the liver was measured using a monoclonal antibody for proteins modified by 4-hydroxynonenal (4HNE). 4HNE is a product of lipid peroxidation and can modify biological molecules through carbonylation. We found significantly higher amounts of 4HNE carbonylation in NASH animals demonstrating a high degree of oxidative stress (Figure 6B).

**Fig 5.**
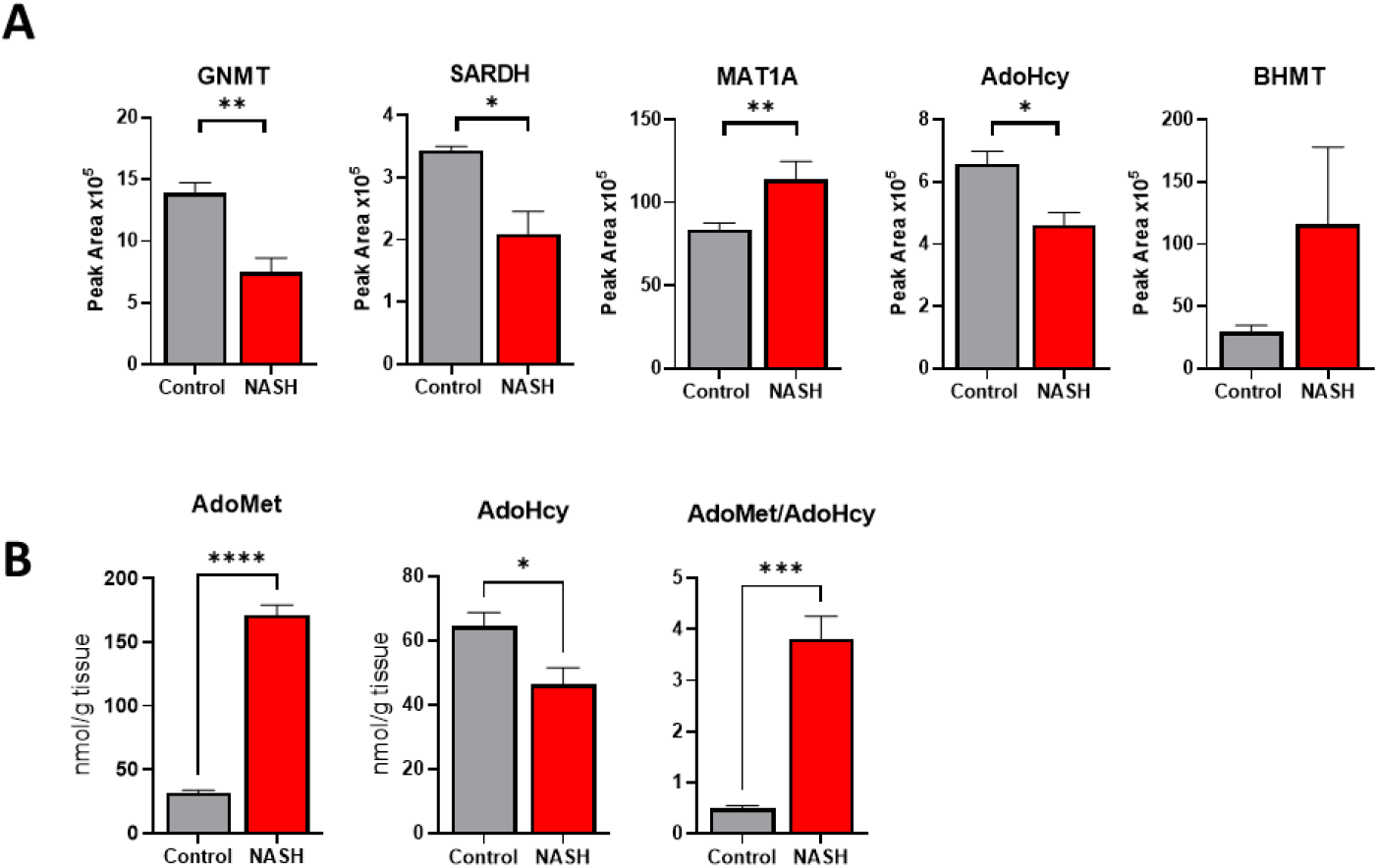
Dysregulation of AdoMet metabolism in NASH. (A) Relative protein abundance of selectAdoMet regulating enzymes in control and NASH livers. (B) Quantification of liver AdoMetand AdoHcy in control and NASH animals. Data presented as mean ± SEM. (N=4 ****p<.0001 ***p<.001 **p<.01*p<.05)

**Fig 6.**
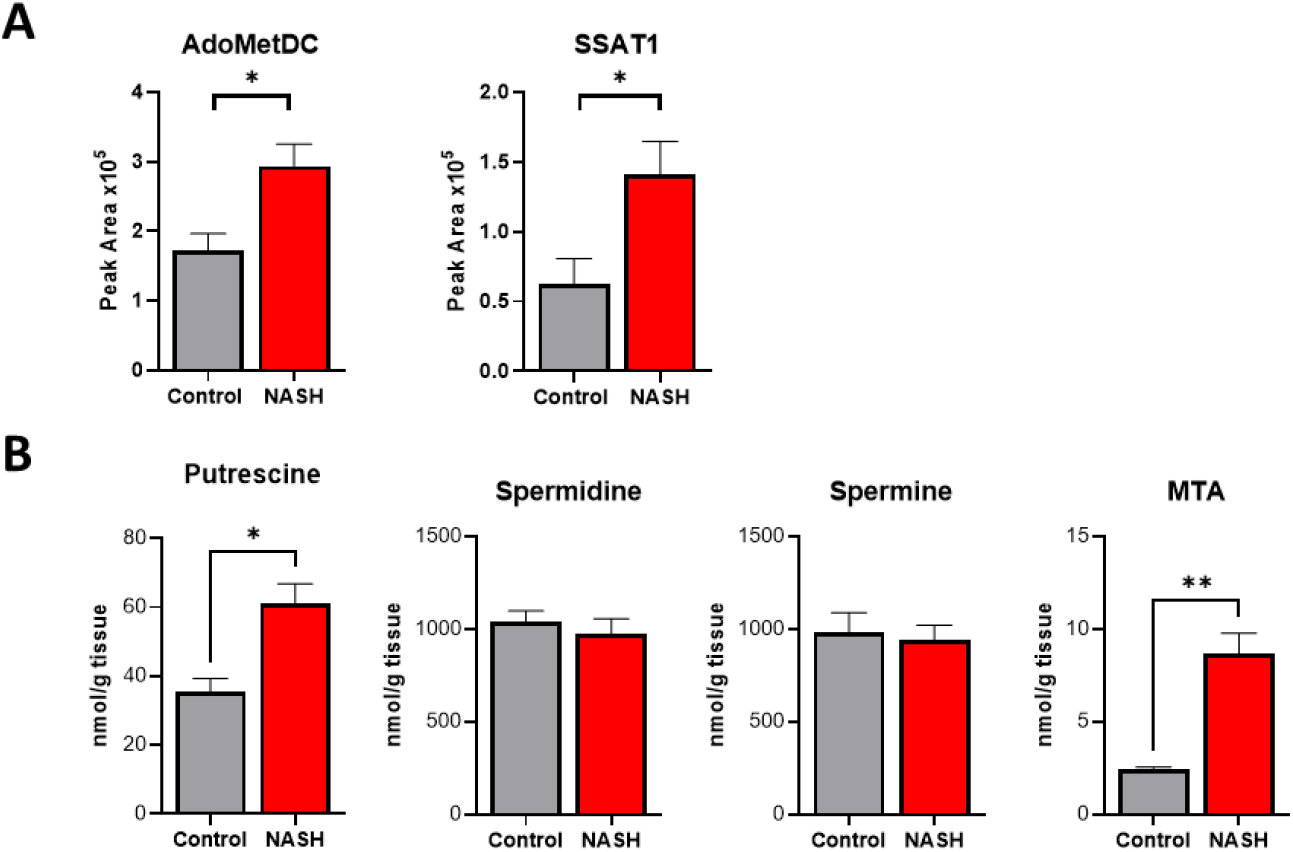
Polyamine metabolism is activated in NASH causing a flux. (A) Relative protein abundance of select polyamine metabolic enzymes in control and NASH livers. (B) quantification of polyamine metabolites in control and NASH livers. Data presented as mean ± SEM. (N=4 **p<.01*p<.05)

**Fig 7.**
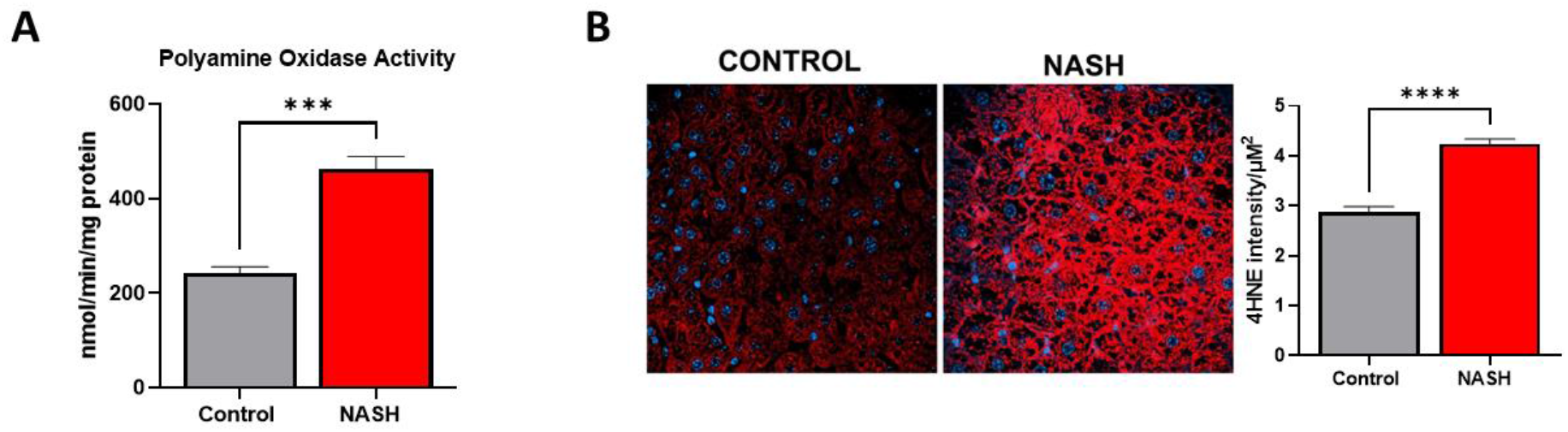
Increased activity of polyamine oxidase and oxidative damage in NASH. (A) Activity of polyamine oxidase in control and NASH livers. (B) lmmunofluorescence staining of control and NASH livers for 4-hydroxynonenal modifiedproteins. Quantification of the fluorescence is shown on the right. Data presented as mean ± SEM. (N=4 ****p<.0001 ***p<.001)

## Discussion

Nonalcoholic steatohepatitis is a largely heterogeneous disease with a complex natural history. One of the major barriers to developing new therapies for NASH is the lack of clinically relevant animal models. Over 70% of NASH patients are obese or have type II diabetes. Using a Western diet-based murine model, we successfully replicated key features of human NASH associated with metabolic syndrome. The animals developed steatosis, lobular inflammation, and fibrosis of the liver while gaining a significant amount of weight and becoming insulin resistant. Our global proteomic analysis provided an in-depth molecular characterization of the disease state and identified key biomarkers and pathways associated with human NASH pathogenesis, including the downregulation of GNMT. We verified the reduction of GNMT in human NASH liver samples and Ingenuity Pathway Analysis identified GNMT downregulation as a key driver in steatohepatitis. We then used targeted mass spectrometry to understand how GNMT downregulation was affecting AdoMet regulation in our NASH model. Our findings correlated well with previous studies using GNMT KO animals showing elevated levels of AdoMet (Martínez-Chantar *et al*., 2008).

Interestingly, the depletion of AdoMet through knockout of synthetic enzyme methionine adenosyltransferase 1 A (MAT1A) or a methionine-choline deficient (MCD) diet has also been shown to lead to NASH (Cano et al., 2011; Itagaki et al., 2013; Lu et al., 2001). Multiple studies have reported deficient AdoMet levels in association with NASH and evidence supporting the supplementation of AdoMet for treatment of NASH (Anstee and Day, 2012; Kalhan et al., 2011; Noureddin et al., 2015). Therefore, the progression of NASH is viable in the presence of either elevated or reduced levels of AdoMet (Jacobs et al., 2013). One explanation for these conflicting findings could be the models used for studying NASH pathogenesis and their translation to human disease. The studies supporting the notion of AdoMet supplementation for NASH treatment rely heavily on data based on the MCD diet. Methionine deficiency leads to decreased levels of AdoMet and results in hypomethylation of DNA and reduced VLDL assembly and export (Caballero et al., 2010; Cano *et al*., 2011; Walker et al., 2011). In our study, we chose to use a Western diet to induce NASH based on the close association of NASH with metabolic syndrome (Younossi and Henry, 2019). The Western diet co-presents with obesity and insulin resistance, whereas the MCD diet induces weight loss. In addition, a recent study comparing the MCD and western diet concluded the human NASH metabolic profile was better represented by the western diet (Machado *et al*., 2015). Therefore, we hypothesize our findings using the Western diet more closely resembles the pathophysiology seen in NASH patients. This is further supported by our findings of reduced GNMT expression in patients with NASH and type II diabetes. The heterogeneity of NASH could explain another possibility for these differences. The progression of NAFLD to NASH is a complex process that results from several genetic and environmental factors (Buzzetti *et al*., 2016). Genetic studies analyzing NASH have identified subgroups of NASH patients with distinct genetic signatures. In a study performed by Alonso et al, they identified an “M subgroup” of NASH patients with a similar genetic signature to MAT1A KO mice. However, this subgroup accounted for less than half of the study population (Alonso et al., 2017). This study illustrates NASH heterogeneity and demonstrates that the MCD diet does not properly model the majority of NASH patients. In addition, a recent meta-analysis of seven human NAFLD genetic data sets identified GNMT downregulation in a genetic signature of NAFLD progression, while MAT1A was not acknowledged (Ryaboshapkina and Hammar, 2017). Our study provides evidence the pathogenesis of NASH in association with metabolic syndrome causes an increase in AdoMet. These findings are further supported by previous studies showing a high-fat diet can lead to increased AdoMet levels in association with NASH. In a study done by Maria Del Bas et al, they use a high-fat diet with selenium and vitamin e deficiency to induce NASH in hamsters (Maria Del Bas et al., 2019). They found AdoMet is significantly upregulated in NASH associated with metabolic syndrome and not altered in simple steatosis. These findings can help stratify the highly heterogenous NASH population into subgroups based on their metabolic profile. The ability to personalize treatment for NASH will greatly help the success of phase three trials and the development of new therapies for NASH.

From our results, a consequence of elevated AdoMet levels was the activation of polyamine metabolism in NASH. AdoMet is a required substrate for polyamine synthesis, and high levels of AdoMet can cause a flux into polyamine metabolism. Both our proteomic and metabolomic analyses demonstrated the activation of polyamine metabolism in NASH, showing increased expression and activity of metabolic enzymes and both polyamine precursors and products of catabolism. These findings are supported by previous studies characterizing GNMT KO mice. Hughey et al. showed the knockout of GNMT caused an increase in several polyamine metabolites to reduce elevated AdoMet levels (Hughey *et al*., 2018). From our understanding, this is the first time polyamine flux has been shown in diet-induced NASH pathogenesis. Intracellular concentrations of polyamines can reach millimolar levels, and catabolism through polyamine oxidase can produce an overwhelming amount of hydrogen peroxide and lipid peroxidation products (Murray Stewart et al., 2018). Oxidative stress, specifically lipid peroxidation, is strongly associated with NASH and is a driver of NAFLD progression. Increased oxidative stress can lead to mitochondrial dysfunction, endoplasmic reticulum stress, and inflammation in NASH development (Koliaki et al., 2015). In this study, we show lipid peroxidation stress marker, 4HNE, was significantly increased in our NASH model. These findings highlight the contribution of polyamine metabolism to the production of oxidative stress. Overall, this study shows the reduction of GNMT in NASH pathophysiology results in an accumulation of AdoMet and polyamine flux leading to oxidative stress production.

## Methods

### Human liver samples

Human liver tissue was purchased commercially from XenoTech. Liver tissue was supplied as frozen prelysate in buffer.

### Animal studies

All animal experiments were approved by the Institutional Animal Care and Use Committee (IACUC). Specific pathogen-free, 6-week-old, male, C57BL/J6 mice of approximately 25 grams body weight were purchased from Jackson’s laboratories, Bar Harbor, ME. Seven days of arrival the animals were weighed, and ear tagged for individual identification. Animals were randomly assigned in two groups (n=7 per group), with no statistical difference in body weight. At 7 weeks of age, the NASH group started receiving high fat/high sucrose diet (Research Diets # D12331i) and at 28 weeks of age the diet was switched to high fat/high fructose/high cholesterol diet (Research Diets # D09100310i). Control animals received CHOW diet. Body weight was monitored weekly. A glucose tolerance test was performed at week 65. Animals were fasted for 5 hours and administered a bolus of glucose (1g/kg) by oral gavage. Blood samples were collected from the tip of the tail and blood glucose was measured at 0, 15, 30, 45, 60, and 120 minutes after the glucose administration. Animals were euthanized at 66 weeks of age and liver tissue was stored at -80°C.

### Histology and NAS scoring

Formalin-fixed liver tissues were paraffin-embedded and sectioned onto slides. Slides from each animal were stained with hematoxylin & eosin (H&E), Mason trichrome, and Sirius red staining. All images were taken at 40x magnification. H&E and Sirius red staining were used to quantitate the NAFLD Activity Score (NAS) and Fibrosis score, respectively. The scores were quantitated based on the scale developed by the Non-alcoholic Steatohepatitis Clinical Research Network. The parameters evaluated were steatosis, lobular inflammation hepatic ballooning and fibrosis (Kleiner et al., 2005).

### Global mass spectrometry analysis

For label-free global proteomics studies the proteins were extracted by adding 6M guanidium hydrochloride buffer and dilution buffer (25mM Tris, 10% acetonitrile). The proteins were digested with Lys-C for 4 hours at 37 ^°^C. Second digestion was achieved by overnight incubation with trypsin. The incubated solution was acidified and centrifuged at 4,500 g for 5 minutes. The supernatants consisting of peptides were loaded onto activated in-house-made cation stage tips (Molina-Franky *et al*., 2021; Zhang *et al*., 2018). The peptides from each sample were eluted into six fractions using elution buffers as previously described (Chen *et al*., 2020; McBrearty *et al*., 2021; Molina-Franky *et al*., 2021; Zhang *et al*., 2018). Mass spec analyses were performed on these fractions using the Q Exactive mass spectrometer (ThermoFisher Scientific). The desalted tryptic peptide samples were loaded onto an Acclaim PepMap 100 pre-column (75 μm x 2cm, ThermoFisher Scientific) and separated by Easy-Spray PepMap RSLC C18 column with an emitter (2 μm particle size, 15 cm x 50 μm ID, ThermoFisher Scientific) by an Easy nLC system with Easy Spray Source (ThermoFisher Scientific). To elute the peptides, a mobile-phase gradient is run using an increasing concentration of acetonitrile. Peptides were loaded in buffer A (0.1% (v/v) formic acid) and eluted with a nonlinear 145-min gradient as follows: 0–25% buffer B (15% (v/v) of 0.1% formic acid and 85% (v/v) of acetonitrile) for 80 min, 25-40% B for 20 min, 40-60% B for 20 min and 60-100% B for 10 min. The column was then washed with 100% buffer B for 5 min, 50% buffer B for 5 min, and re-equilibrated with buffer A for 5 min. The flow rate was maintained at 300 nl/min.

Electron spray ionization was delivered at a spray voltage of 1500 V. MS/MS fragmentation was performed on the five most abundant ions in each spectrum using collision-induced dissociation with dynamic exclusion (excluded for 10.0 s after one spectrum), with automatic switching between MS and MS/MS modes. The complete system was entirely controlled by Xcalibur software. Mass spectra processing was performed with Proteome Discoverer v2.5. The generated de-isotoped peak list was submitted to an in-house Mascot server 2.2.07 for searching against the Swiss-Prot database (Release 2013_01, version 56.6, 538849 sequences) and Sequest HT database. Both Mascot and Sequest search parameters were set as follows: species, homo sapiens; enzyme, trypsin with maximal two missed cleavage; fixed modification, cysteine carboxymethylation; 10 ppm mass tolerance for precursor peptide ions; 0.02 Da tolerance for MS/MS fragment ions.

### Bioinformatic analysis

Differentially expressed proteins from the proteomic analysis were further analyzed using Qiagen’s Ingenuity Pathway Analysis (IPA). Differentially expressed proteins were uploaded to IPA and compared against Ingenuity Knowledge Base reference set. Overrepresented canonical pathways, diseases and functions, and biological networks were identified and Fisher’s exact test was used to determine significance.

### Polyamine metabolomic analysis

Polyamine metabolites were measured by HPLC using the AccQ.Fluor kit (Waters, Milford, Mass.) as described previously (Merali and Clarkson, 1996). Polyamine oxidase activity was measured using soluble protein extracts of mouse liver tissue as previously described (Kramer *et al*., 2008).

### Targeted mass spectrometry analysis

Liver tissue samples were prepared as described in global mass spectrometry analysis. Protein quantification was performed on a TSQ Quantum Ultra triple quadrupole mass spectrometer (Thermo Scientific) equipped with an Ultimate 3000 RSLCnano system with autosampler (Thermo Scientific). The mobile phase consisted of Buffer A, 0.1% formic acid and Buffer B, 85% acetonitrile with 0.1% formic acid. Peptides were separated on an Acclaim PepMap RSLC C18 precolumn (3µm, 100Å) and column (2µm, 100 Å). They were then eluted with a flow gradient of 2-95% Buffer B over 20 minutes. Peptides entered the mass spectrometer through nanoelectrospray ionization with a voltage of 1600 V and capillary temperature 270 °C. The mass spectrometer was run in selected reaction monitoring (SRM) mode detecting the transitions listed in Table S2. Each protein was identified by at least two unique peptides. The selection of peptides for the SRM assay were based on the following: 1. Peptides uniquely representing the target protein 2. No missed tryptic cleavage sites 3. Peptide sequences with 6-25 amino acids 4. Amino acids susceptible to chemical modifications (Cys, Met) avoided. Peptide transitions were identified based on elution profile and fragmentation pattern compared to a spectral library (dotp) generated from our global proteomic analysis or the deep neural network, Prosit (Gessulat et al., 2019). Each peptide was identified by the co-elution of at least 5 product ions. The reproducibility and linearity of each transition was also assessed in a set of replicate liver digests (Figure S1-2). The peptide with the smallest coefficient of variation and greatest coefficient of determination was used for quantification. All peptides used for quantification had a coefficient of variation <20% and demonstrated linearity from 10-200µg of total protein. Data was collected using Xcalibur software and imported into Skyline software for peak area integration and data analysis.

### 4HNE immunofluorescence

Liver sections were subjected to immunofluorescence to evaluate 4HNE-modified protein expression. Briefly, slides were deparaffinized and epitopes were retrieved with citric acid solution followed by incubation in glycine solution. Non-specific background was blocked using 10% goat serum. Slides were incubated with anti-4HNE monoclonal antibody (Abcam ab46545), washed and incubated again with goat anti-rabbit antibody (Thermo Fisher A27034). Nuclei were stained with DAPI. Pictures were taken using confocal microscopy and 4HNE intensity was quantitated with Harmony® software.

### Statistical analysis

Data are expressed as mean values ± standard error of the mean (SEM). Statistical comparisons were performed using unpaired, two-tailed Student’s t tests. Differences were considered significant when P< 0.05.

