## Supplemental Figures 1-3 for "Role of glycine-*N*-methyl transferase in human and murine nonalcoholic steatohepatitis"

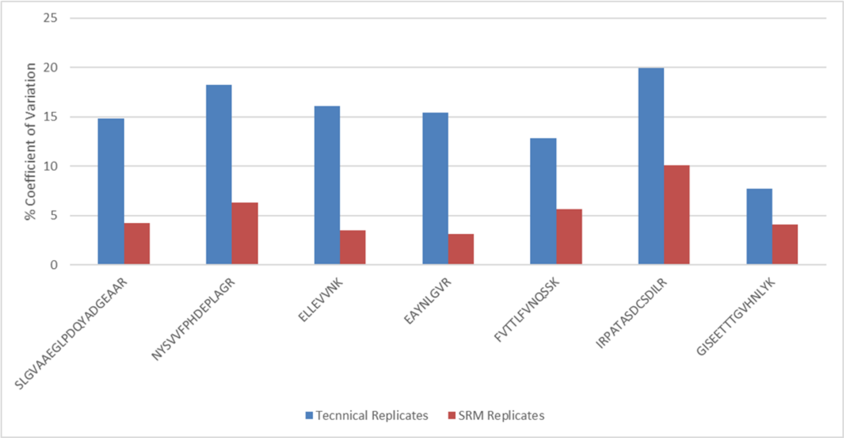


**Supplementary Figure 1.** Five replicate preparations of liver protein lysate were prepared to measure the technical replicate coefficient of variation for each peptide. The SRM replicate variation was assessed by three replicate injections of the same sample preparation. All peptides used for quantification had a coefficient of variation below 20%.

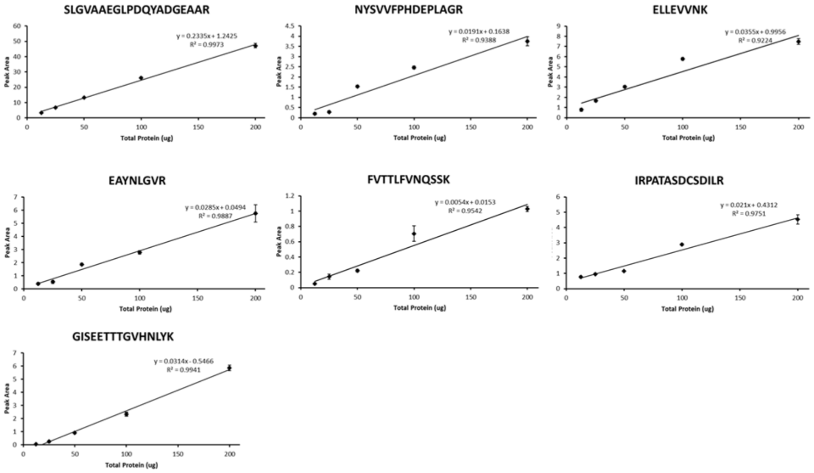


**Supplementary Figure 2.** The linearity of each peptide assay was assessed by preparing liver protein lysate samples with varying amounts of total protein amount and measuring peptide transitions. Samples were prepared in triplicate. The linear regression for each peptide is shown and a total of 100ug of protein was used for subsequent analyses. Data presented as mean ± SEM.

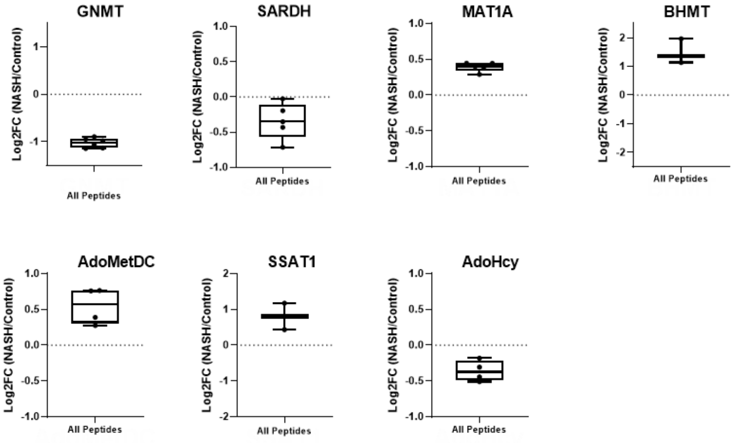


**Supplementary Figure 3.** Box plots showing the log2 fold change for all peptides used for the identification of each protein. The peptide with the lowest coefficient of variation and highest coefficient of determination was used for protein quantification.
